# A community driven GWAS summary statistics standard

**DOI:** 10.1101/2022.07.15.500230

**Authors:** James Hayhurst, Annalisa Buniello, Laura Harris, Abayomi Mosaku, Christopher Chang, Christopher R. Gignoux, Konstantinos Hatzikotoulas, Mohd Anisul Karim, Samuel A. Lambert, Matt Lyon, Aoife McMahon, Yukinori Okada, Nicola Pirastu, N. William Rayner, Jeremy Schwartzentruber, Robert Vaughan, Shefali Verma, Steven P. Wilder, Fiona Cunningham, Lucia Hindorff, Ken Wiley, Helen Parkinson, Inês Barroso

## Abstract

Summary statistics from genome-wide association studies (GWAS) represent a huge potential for research. A challenge for researchers in this field is the access and sharing of summary statistics data due to a lack of standards for the data content and file format. For this reason, the GWAS Catalog hosted a series of meetings in 2021 with summary statistics stakeholders to guide the development of a standard format. The key requirements from the stakeholders were for a standard that contained key data elements to be able to support a wide range of data analyses, required low bioinformatics skills for file access and generation, to have easily accessible metadata, and unambiguous and interoperable data. Here, we define the specifications for the first version of the GWAS-SSF format, which was developed to meet the requirements discussed with the community. GWAS-SSF consists of a tab-separated data file with well-defined fields and an accompanying metadata file.

## Introduction

Summary statistics are defined as the aggregate p-values and association data for every variant analysed in a genome-wide association study (GWAS). The depth of information contained in the summary statistics represents huge potential to extend the power of GWAS and improve disease understanding. In recent years a number of methods have been developed to enable the use of GWAS summary statistics to gain insights into the mechanisms of complex human disease, identify new drug targets and evaluate disease risk. Example methods include large meta-analyses (Wheeler E. 2017), trait pleiotropy (Smeland O.B, 2017), prediction using polygenic scores (PGS) (Lambert S, 2019) and Mendelian randomisation (MR) (Paternoster L, 2017). However, a considerable number of summary statistics are still not fully and openly shared with the community, either being made available under controlled access, upon agreement to restrictive terms, with incomplete data, or not shared at all. One of the main challenges associated with sharing full GWAS results is the lack of standards for data content and format, meaning that researchers do not have clear guidelines for appropriate file generation for sharing, and the re-usability of the resulting files can be poor. Typically, each GWAS will produce a single file with a table of summary statistics containing a list of variants with p-values, other statistics and relevant annotations or metadata. Generated by different software packages and made available via different resources, summary statistics can vary in a myriad of ways from one study to the next. A recent analysis of 327 summary statistics files found over 100 unique formats (Murphy et al, 2021). Differences in file formats, header definitions, data types, genetic variant or association data reporting and missing data create challenges for users by reducing data interoperability.

The GWAS Catalog began hosting summary statistics in 2018, and rapidly developed a first minimal data format based on the most commonly included fields in publicly available files (Buniello, 2019), but without community input. In parallel, other summary statistics formats have been defined for specific purposes, e.g. dbGap’s Minimum Information Required for Association Data guidelines, designed to fulfil data sharing requirements in dbGaP (https://www.ncbi.nlm.nih.gov/gap/docs/submissionguide/#apha); GWAS-VCF (Lyon, 2021) developed for robustness and performance and to underpin the OpenGWAS platform (Elsworth, 2020) and associated tools. However, key data fields to support the widest range of downstream uses, and mandatory metadata content, have not previously been defined. The field is advancing rapidly and summary statistics data sharing is quickly becoming more common. More than 70% of GWAS Catalog studies are now linked to freely accessible summary statistics (27,500 studies from 550 publications) with the highest yearly increment observed in 2020-2021, and 77% of summary statistics submissions to the GWAS Catalog in 2021 were made before publication, upon a journal’s mandate. These metrics show that human GWAS summary statistics have now reached a critical mass, and to maximise the utility of this body of data there is a need for the community to adopt an information standard to which users can expect all studies to adhere (MacArthur et al, 2021). A single standard with stricter definitions on the data included will increase the utility of GWAS summary statistics, reduce the risk of misinterpretation of data and enable users to easily analyse and integrate data from different GWAS. A range of mandatory data fields are required to support the major use cases for downstream analysis, such as PGS development, MR, meta-analysis and functional annotation of variants, at scale.

## Methods

Following initial discussions with the GWAS community at the 2020 GWAS summary statistics standards and sharing workshop (MacArthur et al, 2021), the GWAS Catalog hosted a series of meetings between June 2021 and September 2021 with invited summary statistics stakeholders including data generators, data users, data managers and bioinformaticians, representing diverse user groups. These meetings gathered requirements and identified challenges. The aim of this process was to finalise minimum information elements for data sharing to maximise downstream utility, and to complete a phase of iteration on the proposed standard.

An initial set of standard reporting elements had been proposed in MacArthur et al, based on a public survey and workshop discussion. The first meeting reviewed these findings, assessed currently available formats with their strengths and weaknesses, and then comprised a guided discussion focusing on the details of a potential new format. Topics covered included user considerations, in particular the balance of requirements for data consumers and data generators; data reporting requirements including consistent and unambiguous variant reporting and p-value precision; mandatory and optional fields for data and metadata; storage and access of metadata; scaling considerations. Following the first meeting we circulated a survey amongst meeting attendees to summarise opinions on variant reporting, association reporting, metadata fields and location, and file format. The outcome of the survey was reviewed and discussed at the second meeting with the goal of reaching consensus on requirements. Based on the identified requirements, GWAS Catalog staff developed a proposed format which was presented and iterated at the third meeting in Sept 2021. Wider feedback was gathered from the community via github and email between the initial preprint release of this manuscript in July 2022 and March 2023.

### Impact assessment

In order to evaluate the potential impact of the proposed format on data sharing, we assessed the content of existing submissions to the GWAS Catalog. To date, the minimum requirements for submission are a p-value with either rsID or chromosome and base-pair location, with other fields supplied at the submitter’s discretion.

Taking all author-submitted single-variant GWAS summary statistics files in the GWAS Catalog (315 submissions, 27845 GWAS) as a sample, we used GNU grep to search all the files for the newly defined mandatory fields from the proposed format. We searched the first row of each file for a) the presence of each mandatory field; and b) the presence of all the mandatory fields (an OR operator was used between beta and odds_ratio).

## Results

### Requirements

The key requirements for an expanded GWAS summary statistics standard obtained from the stakeholders’ use cases were as follows:

- Consistent representation of data to enable interoperability
- Easily accessible metadata for summary statistics to facilitate data interpretation and re-usability
- Unambiguously reported genetic variants for standard annotation
- A set of mandatory (i.e. must be present and filled with non-null values) fields, providing the information necessary to enable a wide range of data analyses including MR and PGS development
- A set of encouraged fields with standard headers, which are strongly recommended but not mandatory
- A balance between these mandatory and encouraged fields that includes essential data but does not set the bar impossibly high for the community using and implementing the standard
- A low bioinformatics requirement for data consumers and data producers, reflecting the composition of the user community, to maximise stakeholder uptake
- The format should be interoperable with other major formats and resources.

These requirements were used to define the backbone of a format - the GWAS-SSF. We solicited public feedback on the proposed format via our github repository https://github.com/EBISPOT/gwas-summary-statistics-standard and via email, and made iterative changes in response, including the addition of a metadata field for sex and consideration of the reference allele. The format was fully implemented in the GWAS Catalog submission, data release and harmonisation pipelines and as an output option in PLINK 2.0 in April 2023.

## GWAS-SSF, a newly proposed GWAS summary statistics format

The GWAS summary statistics format (GWAS-SSF) is composed of two data components, the summary statistics data and accompanying metadata.

### Summary statistics data format

The GWAS-SSF data file is a TSV flat file of tab-delimited values that can be compressed (see Figure 1 for a schematic representation, Supplement 1 for example file), reporting data from a single genome-wide analysis. The first line of the file contains the headers to the table. The rows after the header store the variant association data. Where permitted, values can be omitted by the presence of “#NA”. There are no limits to the number of rows or columns that the table can have, however, a set of mandatory fields (defined in Table 1) must be present in a defined order. A file may contain additional columns beyond the set of mandatory fields. Table 1 shows some non-mandatory (encouraged) fields that may be present.

**Figure 1.**
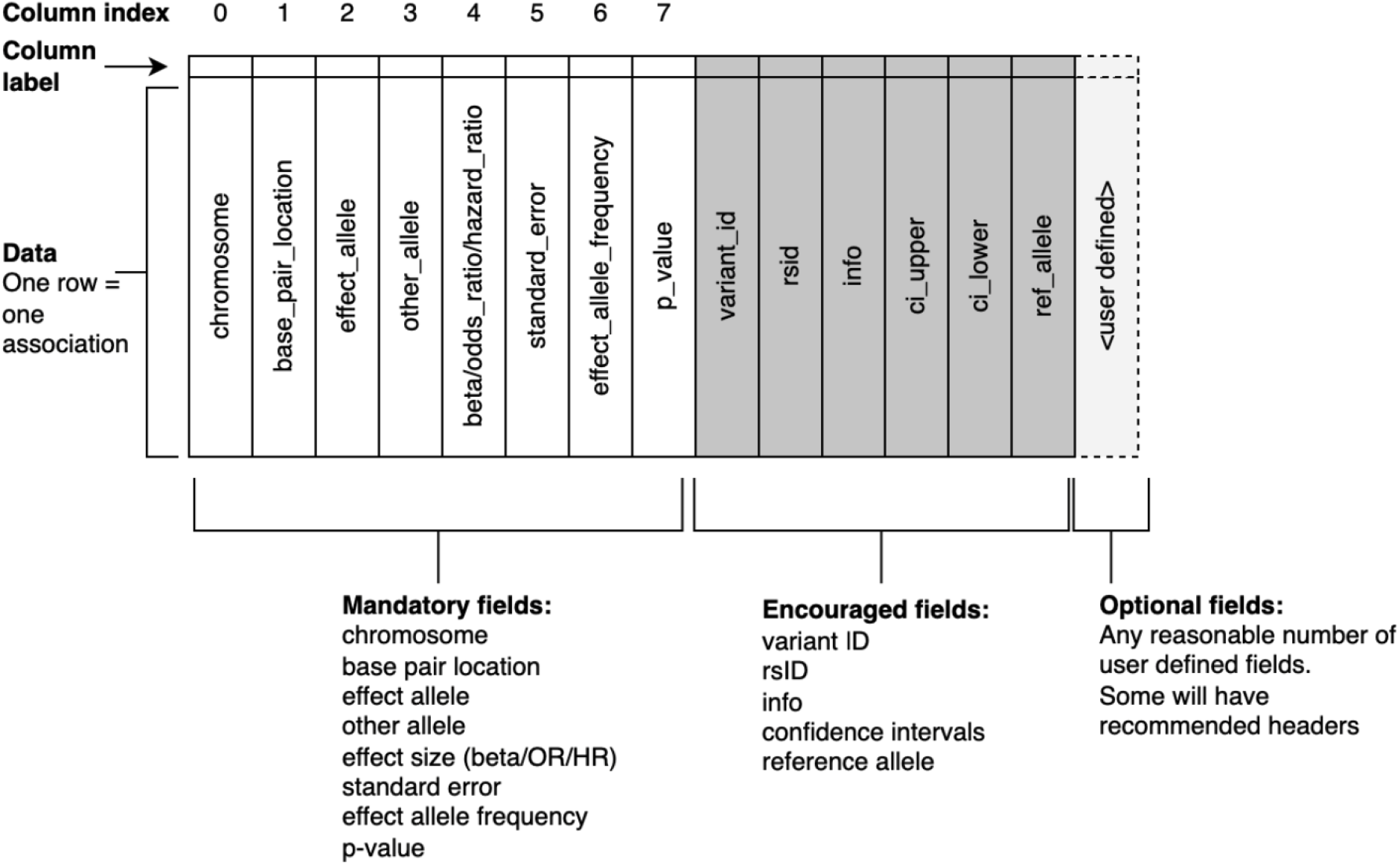
Schematic representation of the summary statistics table. Examples of data content within each specific field are provided in Table 1.

**Table 1.**
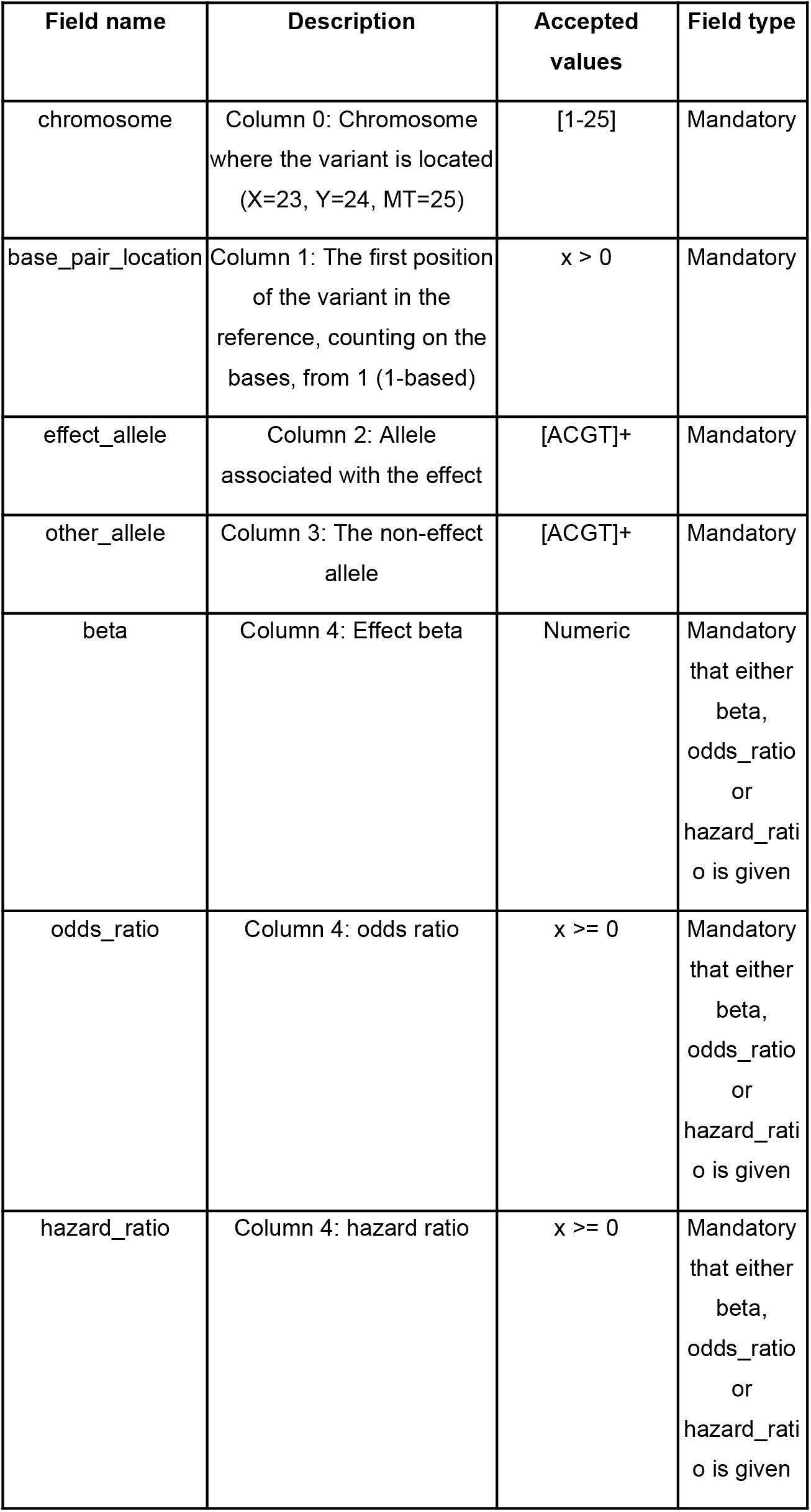
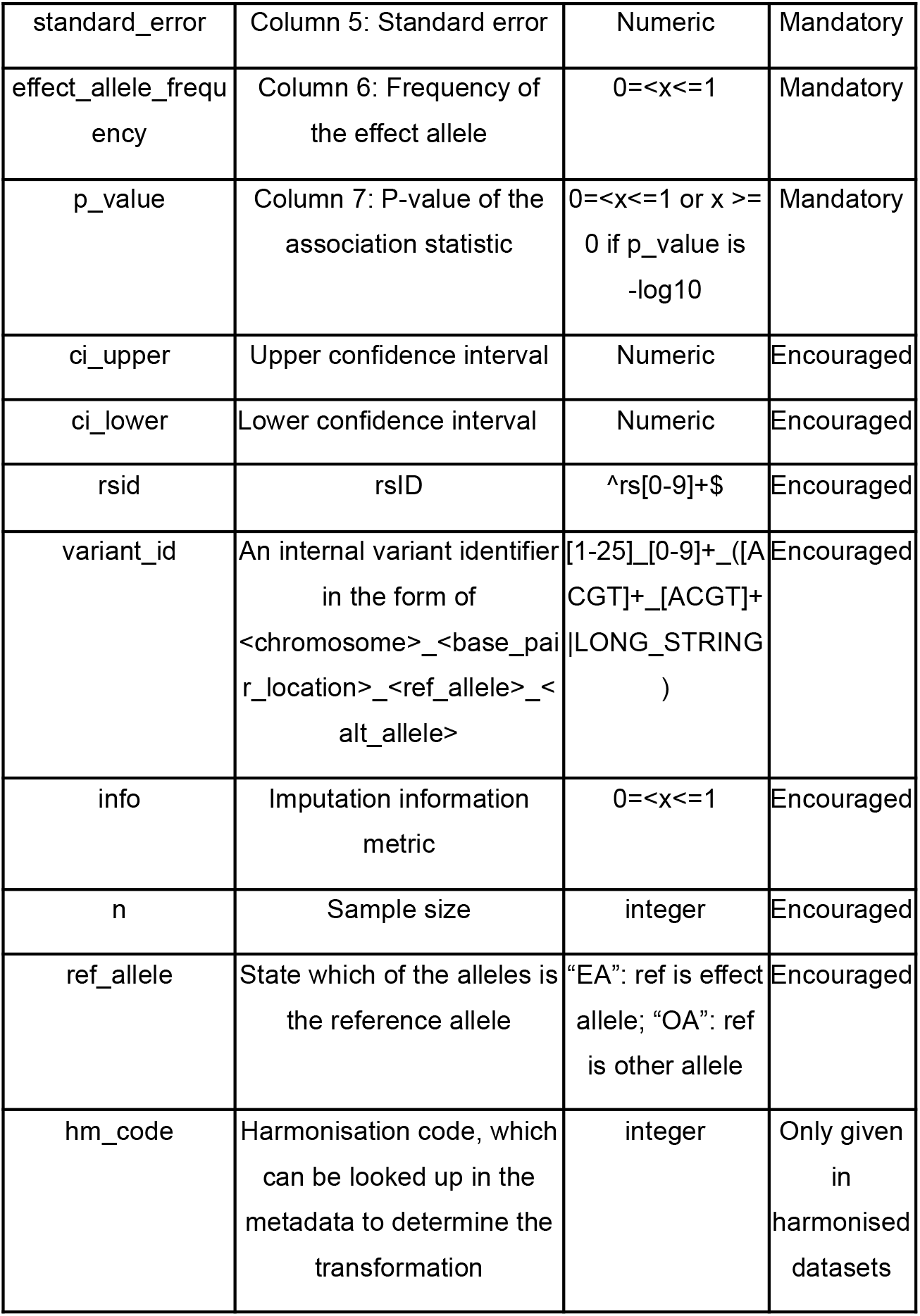
Summary statistics field definitions. Note 1: If p-value is equal to 0, the precision of the p-value calculation must be given in the accompanying metadata. Note 2: Effect allele frequency can be rounded up to a threshold value defined in the metadata.

### Summary statistics table contents

Four fields in the summary statistics table, combined with the reference genome assembly provided in a metadata file (see below), define the genetic variants (all field definitions can be found in Table 1). These fields are the chromosome (*chromosome),* the genomic location position on the chromosome (*base_pair_location*), the effect allele (*effect_allele*), and the non-effect allele (*other_allele*). Chromosome values are integers from 1 to 25, with chromosome X mapping to 23, chromosome Y to 24, and mitochondrial to 25. Genomic location is an integer value representing the first position of the variant in the reference genome, using the coordinate system specified in the metadata, either 1-based or 0-based (see Figure 2). The *effect_allele* field captures the allele for which the effect is associated, while the *other_allele* field reports the non-effect allele. Both of the allele fields will contain allele strings, including cases where variants are insertions and deletions (see Figure 2). *variant_ID* is for storing *chromosome, base_pair_location,* reference and alternate allele information as a single concatenated (with underscores) string and rsID can be stored in the *rsid* field, but both fields are optional.

**Figure 2.**
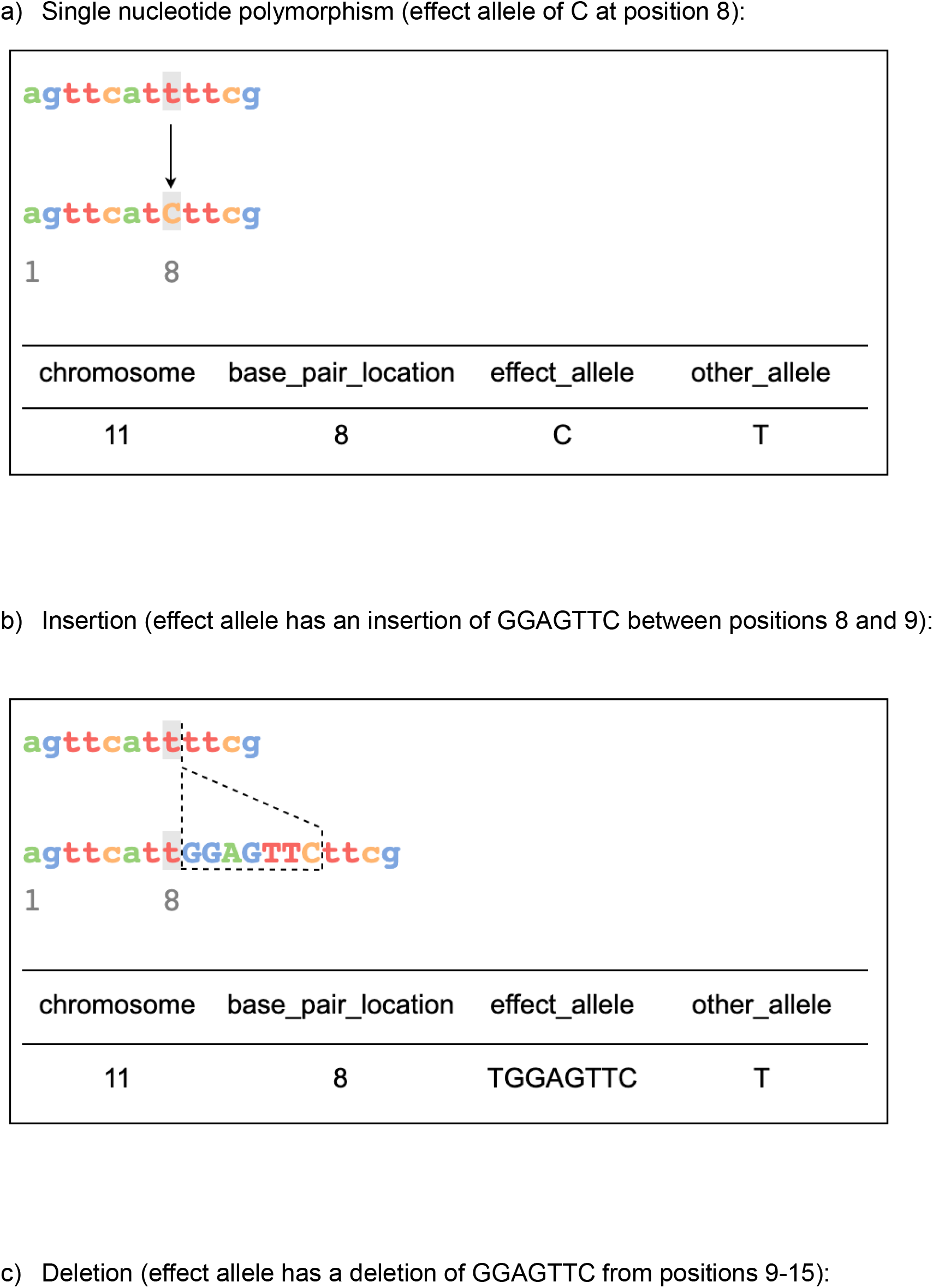
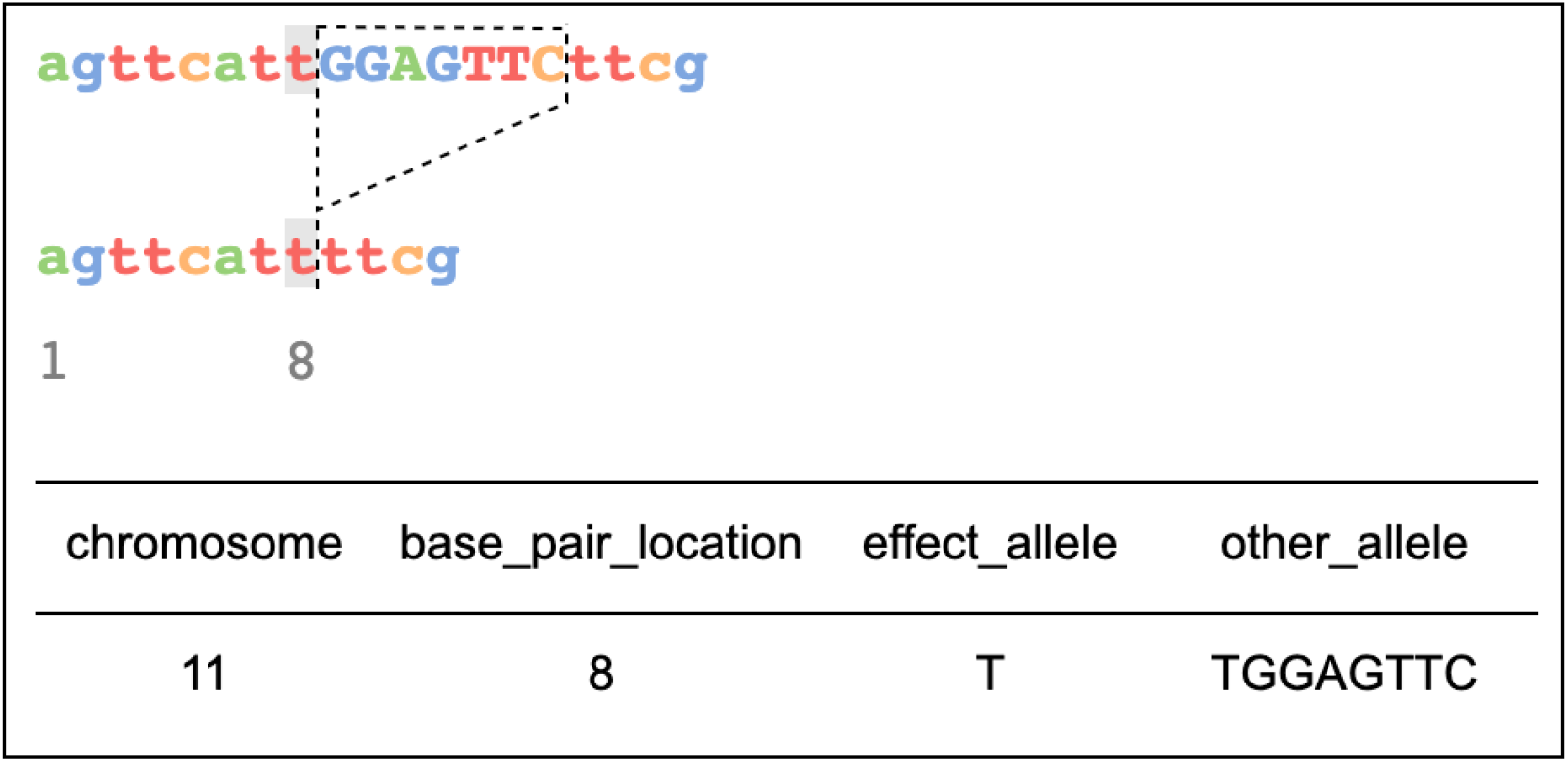
Illustration of how variants are recorded in the summary statistics table for (a) SNP, (b) insertion, and (c) deletion alleles. Note that for insertions and deletions, the position of the base preceding the indel (the highlighted T at 8) is the position used to index the variant.

For optimum reusability of files, genomic location should be represented consistently using the same coordinate system. However, GWAS-SSF is designed to be flexible in order to promote data sharing, even of legacy data, with minimal effort. The coordinate system is indicated in an optional metadata field. For downstream use, the GWAS Catalog provides a harmonised file (see below) that is aligned using 1-based indexing to maximise interoperability with variant call format (VCF) (Danecek et al 2011).

Similarly, the format is flexible with regards to allele orientation. We recommend that analyses be performed with the other allele set to the reference, since indels cannot be unambiguously identified without knowledge of the reference allele, with resulting loss of information. However, many datasets exist with unordered alleles, and the format is flexible to share such data. The field *ref_allele* indicates whether the effect or other allele is the reference. For downstream use, GWAS Catalog harmonised files contain data oriented to the reference strand, i.e. the effect and other alleles are equivalent to the alternate and reference alleles, respectively.

All rows contain the following association statistics: p-value (*p_value or neg_log_10_p_value*), the effect size (either *beta, odds_ratio* or *hazard_ratio*), and the standard error (*standard_error*). Depending on the precision of software that performed the calculation of association, p-values in GWAS analyses may appear rounded to zero or one. This is particularly problematic where highly significant associations (e.g. p<10^-300^) are rounded to zero, preventing associations being ranked in order of significance. Calculation of accurate p-values is recommended where possible. Where this is not possible due to limitations of the software used, the GWAS-SSF requires the analysis and genotype imputation software and version to be present in the metadata, to help users of the summary statistics interpret these values. Alternatively, p-values can be expressed as negative log values, in which case the p-value column header should be *neg_log_10_p_value*. Effect allele frequency (*effect_allele_frequency*) is a mandatory field. However, where privacy concerns might otherwise be a barrier to sharing the data, a cutoff may be specified in the metadata (*minor_allele_freq_lower_limit* field, see Table 2) so that frequencies below that cutoff are rounded-up to mask their true values. For example, *minor_allele_freq_lower_limit* = 0.01 in the metadata file would communicate that the lowest possible value for the minor allele frequency in this file is 0.01, and anything below this threshold has been rounded up to 0.01.

**Table 2.** Metadata field definitions

| Field | Description | Accepted value | Mandatory |
| --- | --- | --- | --- |
| genome_assembly | Genome assembly | GRCh/NCBI/UCS C value | Yes |
| coordinate_system | Coordinate system | 1-based/0-based | Yes |
| trait_description | Author reported trait description | Text string (multiple possible) | Yes |
| sample_size | Sample size | Integer | Yes |
| case_count | Number of cases for case/control study | Integer | No, unless caseControlStudy is true |
| control_count | Number of controls for case/control study | Integer | No, unless caseControlStudy is true |
| case_control_study | Flag whether the study is a case-control study | Boolean | No (default is false) |
| sample_ancestry | Sample ancestry | Text string (multiple possible) | Yes |
| genotyping_technology | Genotyping technology | Text string (multiple possible) | Yes |
| analysis_software | Software and version used for the association analysis | Text string | Yes if p-values of 0 given |
| imputation_panel | Imputation panel | Text string | No |
| imputation_software | Software used for imputation | Text string | No |
| minor_allele_frequency_lower_limit | Lowest possible minor allele frequency | Numeric | No |
| ancestry_method | Method used to determine sample ancestry e.g. | Text string (multiple possible) | No |
|  | self-reported/genetically determined |  |  |
| is_sorted | Flag whether the file is sorted by genomic location | Boolean | Yes |
| adjusted_covariates | Any covariates the GWAS is adjusted for | Text string<br>(multiple possible) | No |
| ontology_mapping | Short form ontology terms describing the trait | Text string<br>(multiple possible) | No |
| sex | Indicate if and how the study was sex-stratified | "M", "F",<br>"combined" | No |

### Summary statistics metadata

Metadata accompanying the data file describes the summary statistics, such as the name and md5sum of the data file (see Supplement 2 for example), and sample and experimental metadata of the GWAS itself (Table 2), thereby ensuring the reusability of the data. The metadata file fields can be expanded as needed in the future, and as with the summary statistics file, additional columns can be included as required. Sample metadata fields include descriptions of the trait under investigation and the sample size and ancestry. An additional field *ancestry_method* can be used to indicate whether the ancestry descriptor is self-reported or genetically defined (encouraged). We recommend that ancestry is reported according to the standardised framework guidelines described in Morales et al, 2018. Every effort should be made to explicitly note whether the sample is admixed and the ancestral backgrounds that contribute to admixture. The trait description is free text and should include a clear description of the trait under study, including any relevant background characteristics of the study population, e.g. “lung cancer in asthma patients”. Trait ontology terms can be stored in the metadata o*ntology_mapping* field. There are both mandatory and encouraged metadata fields, which are detailed in Table 2.

## Implementation in the GWAS Catalog and other resources

### Metadata availability

Each summary statistics file will be made available with an accompanying metadata file in YAML format, “a human-friendly data serialisation language for all programming languages“ (https://yaml.org/). This format aims to strike a compromise between human- and computer-readability and is easily convertible to json. Data and metadata files located in the same folder and are linked by the GWAS Catalog accession ID. Metadata submission is via a structured excel template which is validated and QC’d. Yaml files are generated from the submitted metadata upon data release. Metadata files will be generated retrospectively for all pre-existing summary statistics in the Catalog.

### Support for data submission

Previous discussions with the community suggested that the burden of formatting data should lie with data generators (MacArthur et al, 2021). However, with increasingly large datasets, the effort and associated cost required in preparing files for submission to repositories is a key concern raised by our working group attendees and others (Kozlov 2022), and support is required to mitigate the burden. The GWAS Catalog provides easy to use tools for formatting, converting to the standard headers and checking the validity of summary statistics files prior to submission, requiring input of a simple TSV file (https://github.com/EBISPOT/gwas-sumstats-tools). The validator runs upon submission of summary statistics to the GWAS Catalog, and must pass in order for data to be successfully submitted. An offline version is provided for users to check the validity of their files prior to upload with detailed feedback provided on failures.

PLINK2.0 (www.cog-genomics.org/plink/2.0/; Chang et al 2015), now includes an option--gwas-ssf to generate results files in the GWAS-SSF format, thus removing the need for data generators to further manipulate files after analysis for submission to the GWAS Catalog.

Summary statistics submitted to the GWAS Catalog are accessioned and citable at point of submission, even prior to journal publication.

### Harmonised datasets

In addition to the author-submitted summary statistics that the GWAS Catalog makes available for download, eligible summary statistics are also made available to users in a harmonised format (Buniello, 2019; https://github.com/EBISPOT/gwas-sumstats-harmoniser). The harmonised data have been oriented to the same reference strand so that the harmonised summary statistics of one study are interoperable with any other harmonised study. Newly harmonised files will adhere to the GWAS-SSF format described here but benefit from being sorted and indexed by genomic location and compressed with bgzip, allowing fast retrieval of variants of interest by location.

### Interoperability with other resources

GWAS-SSF was designed to be compatible with GWAS-VCF, and can be converted using publicly available tools. dbGaP will accept submissions of GWAS summary statistics in the same format to ensure flow of data between these two important public resources. We hope that other resources will mandate the standard data content and headers defined in GWAS-SSF to enhance interoperability and maximise the number of usable datasets.

## Impact assessment

Of 27,845 valid summary statistics files obtained by the GWAS Catalog between the release of our submission system in 2020 and July 2022, ∼17% were missing at least one of the new mandatory fields (Fig 3(a)). The most commonly omitted field was effect allele frequency, followed by standard error and effect size (beta/odds_ratio/hazard_ratio). Each submission may contain multiple files, with the number varying from 1 to thousands, and it is reasonable to assume that all the files within a submission adhere to the same format, being generated as part of the same research project. We therefore wished to ascertain the proportion of *submissions* that were missing data, as this may be more indicative of practices within the data-generating community. More than 50% of submissions contained files omitting at least one of the new mandatory fields, with the most commonly omitted field again being effect allele frequency followed by standard error, other allele and effect size (beta/odds_ratio/hazard_ratio) (Fig 3(b)).

**Figure 3:**
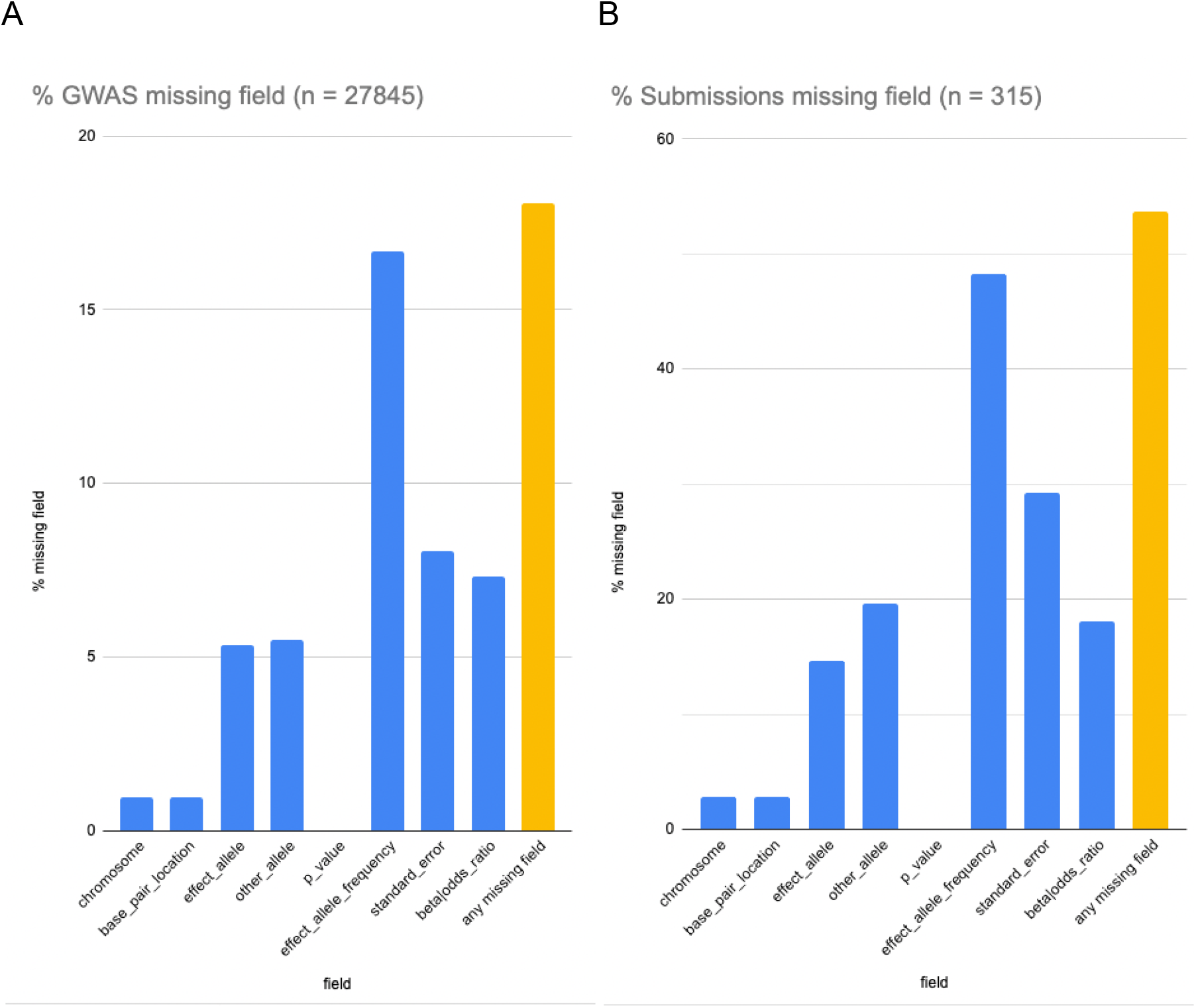
Assessment of missing data in summary statistics shared under minimal requirements, based on (a) individual GWAS datasets and (b) submissions, each typically associated with a single manuscript or project.

These results show that a substantial portion of GWAS summary statistics shared under minimal requirements is severely limited in usability for downstream purposes. In the case of effect size, the summary statistics will be rendered unusable for the majority of methods that leverage summary statistics. Since more than 50% of submissions omitted at least one of the new mandatory fields, implementation of GWAS-SSF has the potential to double the number of usable datasets for important downstream uses.

## Discussion

Community activities have been effective in the development of agreed standards and sharing principles for scientific data (e.g. Brazma et al, 2001). The GWAS summary statistics format (GWAS-SSF) presented here is the result of meetings with the community to make a simple, easy to access standard which promotes cross-dataset consistency and is useful for varied use cases. The mandatory content of the table meets the requirement set by the stakeholders of the working group to perform most analyses e.g. beta and standard error to support MR analyses, effect and non-effect allele to support meta-analysis and generation of polygenic scores. Another requirement from the working groups is a consistent approach to variant reporting and representation which is important for users of the data to be able to easily merge or compare datasets. By adopting the variant reporting standard embodied in VCF (Danecek et al 2011) to define single nucleotide polymorphisms and short indels, consistency and interoperability will be achieved. More complicated variants (e.g. structural variants) and their shorthand notations fall outside the primary scope of GWAS summary statistics. Regarding file type, the GWAS-VCF format (Lyon et al 2021; Elsworth et al 2020) is well suited to high throughput use, but less-technical stakeholders preferred a TSV file with easily human-readable metadata, and comparable files are widely used in the field. We have therefore implemented the standard data content in a generic TSV/YAML format as a commonly used file type for GWAS summary statistics which is universally accessible and can be converted to GWAS-VCF or other high throughput format as required.

In our implementation, each GWAS analysis is represented in a separate file, differing from GWAS-VCF which can represent multiple phenotypes in the same file. Although there are advantages in sharing data between individual users in this way, the number of GWAS per unit is rapidly growing, for example >18K phenotypes in Wang et al 2021, and this may cause usability issues to the average user where large volumes of data are stored in a single file.

In the GWAS Catalog’s implementation, metadata for the summary statistics files and study design are available in a separate file, allowing a generic tabular data file format (TSV) which is universally accessible using readily available tools. Data and metadata files are labelled with their GWAS Catalog accession ID, which in turn links the files to additional metadata, top associations, polygenic scores and other annotation made available via the GWAS Catalog. The metadata file is an optional source to help describe the data and reduce ambiguity, without adding complexity to the data file itself. An alternative strategy is to store metadata within the header of data files, and there are advantages to this, primarily that the metadata and summary statistics cannot become inadvertently decoupled. The community should consider whether this is a necessary future requirement for GWAS data, with appropriate tooling.

GWAS-SSF includes a number of mandatory fields, and we heard from our working group that many more fields may be important in certain contexts, e.g. imputation info for filtering variants to identify those of high enough quality for downstream analyses, such as fine-mapping, enrichment analyses, MR, or genetic correlation estimation. However, there was an acceptance that these may not be readily available or necessary for all users and their absence should not preclude data sharing and reuse. The standard should promote open data sharing as widely as possible, while providing the essential information for most major downstream uses. We have therefore included additional encouraged fields with standard headers to promote interoperability, and data generators are strongly encouraged to share these data unless they are genuinely unavailable (for example, in the case of historical/legacy data) or there is a scientific or ethical reason not to (e.g. privacy concerns). Furthermore, the list of standard fields is not intended to be exhaustive and data generators are encouraged to share as much additional data as possible.

A key point of discussion is whether alignment of the non-effect and effect alleles to the reference and alternate alleles can be enforced at the point of submission. Without alignment of alleles, key information on the identification of indels is lost and thus there is a strong argument to be made for a mandate. However, there are two scenarios to balance here: (i) the scientist has easy access to a VCF file which spells out the REF/ALT alleles for these otherwise-ambiguous indels, and (ii) they do not. In the second scenario, it is better for the scientist to share the results they have than to share nothing at all. However, we urge data generators to consider this at the point of data analysis, especially as more dense imputation panels with better coverage of indels become available. A mandatory requirement for allele alignment should be considered in future iterations.

Lastly, development of the format has been driven by the needs of human genetics researchers, and this is a limitation with respect to potential application of the format to other species. Future development work should focus on cross-species capability, to maximise interoperability.

FAIRification of GWAS results is currently a significant challenge for the human genetics community, as thoroughly discussed in our working group meetings and reported in this work. We have described here a community-driven minimum information standard within a simple and universal format that conforms to the FAIR principles (Wilkinson et al 2016) and has been implemented in the GWAS Catalog to maximise usability and accessibility for all potential users of summary statistics. We recommend that mandatory data elements are adopted as widely as possible.

## Data availability

The GWAS Catalog is an open-source project and code is available in the project’s github repository (https://github.com/EBISPOT/goci). Summary statistics data are available via the GUI https://www.ebi.ac.uk/gwas/downloads/summary-statistics and API (http://www.ebi.ac.uk/gwas/summary-statistics/api/). GWAS summary statistics submitted after March 2021 are made available under CC0 terms (https://creativecommons.org/publicdomain/zero/1.0/), while those submitted prior to March 2021 are made available under the standard terms of use for EBI services (https://www.ebi.ac.uk/about/terms-of-use/). We advise consumers of summary statistics data hosted by the GWAS Catalog to note the licence terms of individual datasets. Other GWAS Catalog data is covered by the EBI terms of use. Code is available under the Apache version 2.0 licence (http://www.apache.org/licenses/LICENSE-2.0).

## Supporting information

Example of summary statistics TSV data file

Example of summary statistics metadata YAML file

## Acknowledgements

Special thanks to Mike Feolo, working group participants and the wider community for their engagement and contributions. Research reported in this publication was supported by the National Human Genome Research Institute of the National Institutes of Health under award no. U41HG007823 and EMBL-EBI Core Funds. In addition, we acknowledge funding from: the European Molecular Biology Laboratory; I.B., “Expanding excellence in England” award from Research England; ML, the MRC Integrative Epidemiology Unit (MC_UU_00011/4), supported by the NIHR Biomedical Research Centre at University Hospitals Bristol and Weston NHS Foundation Trust and the University of Bristol. The content is solely the responsibility of the authors and does not necessarily represent the official views of the National Institutes of Health, the NHS, the National Institute for Health and Care Research or the Department of Health.

---

**Supplement 1.**
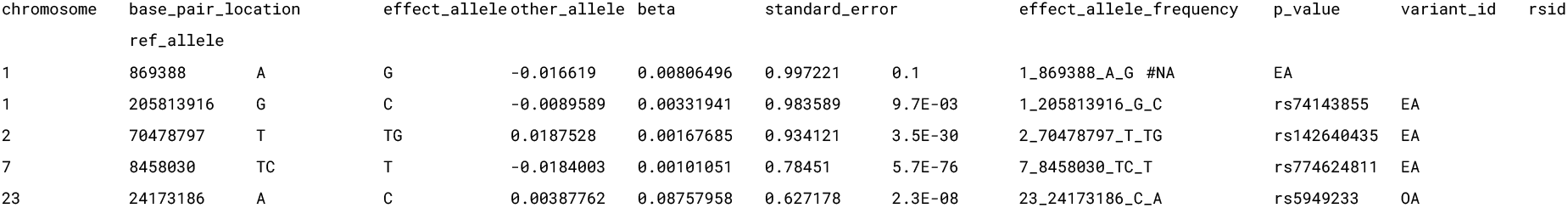
Example of summary statistics TSV data file

**Supplement 2.**
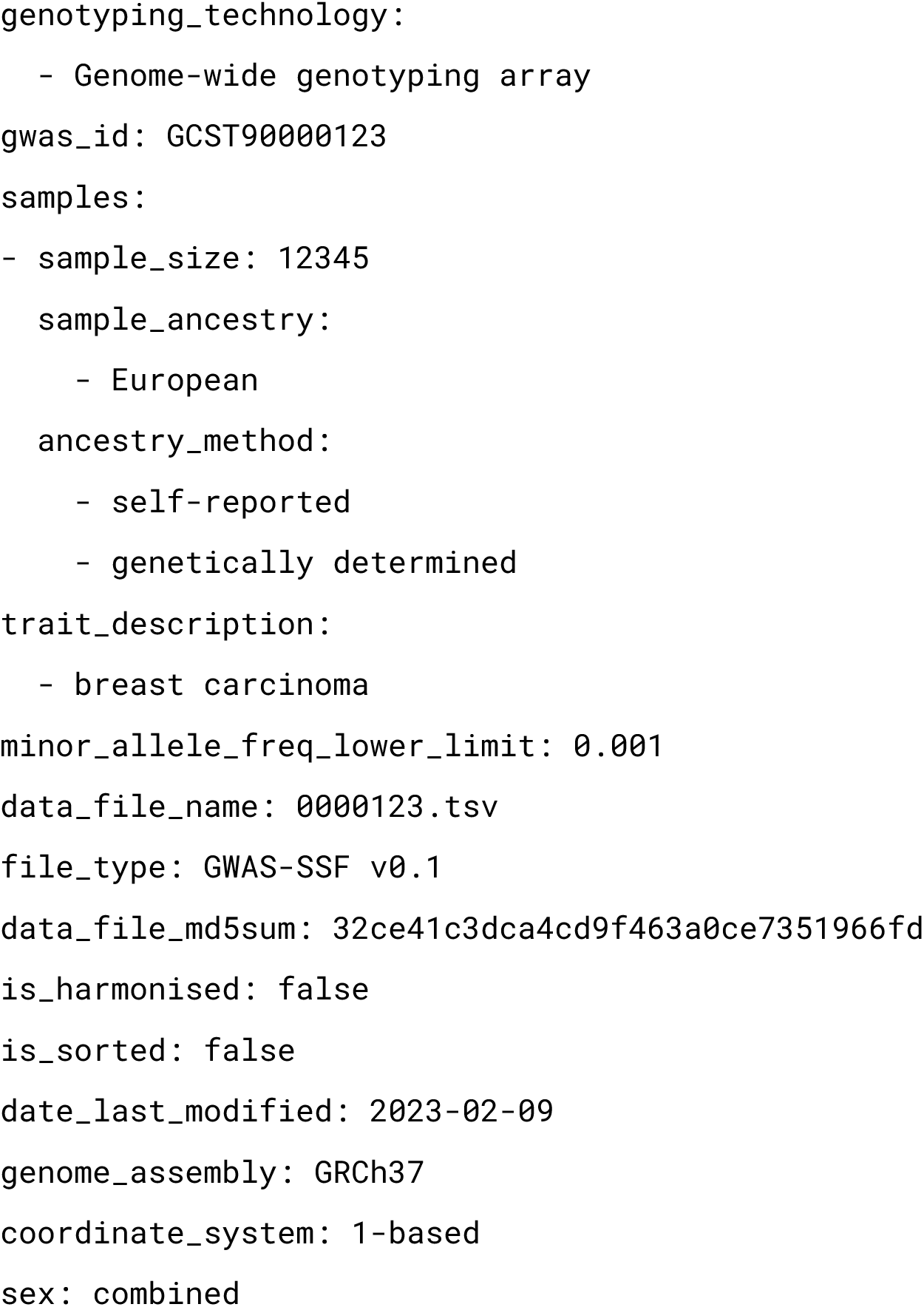
Example of summary statistics metadata YAML file

## Notes

### Competing Interest Statement

The authors have declared no competing interest.

### Summary of Updates

Removal of variant normalisation by left-align and trim from harmonised data description.

